# Disruption of synaptic transmission in the Bed Nucleus of the Stria Terminalis reduces seizure-induced death in DBA/1 mice and alters brainstem E/I balance

**DOI:** 10.1101/2021.12.23.473665

**Authors:** Maya Xia, Benjamin Owen, Jeremy Chiang, Alyssa Levitt, Wen Wei Yan, William P. Nobis

## Abstract

Sudden unexpected death in epilepsy (SUDEP) is the leading cause of death in refractory epilepsy patients. Accumulating evidence from recent human studies and animal models suggests that seizure-related respiratory arrest may be important for initiating cardiorespiratory arrest and death. Prior evidence suggests that apnea onset can coincide with seizure spread to the amygdala and that stimulation of the amygdala can reliably induce apneas in epilepsy patients, potentially implicating amygdalar regions in seizure-related respiratory arrest and subsequent postictal hypoventilation and cardiorespiratory death. This study aimed to determine if an extended amygdalar structure, the dorsal bed nucleus of the stria terminalis (dBNST), is involved in seizure-induced respiratory arrest (S-IRA) and death using DBA/1 mice, a mouse strain which has audiogenic seizures and a high incidence of postictal respiratory arrest and death. The presence of S-IRA significantly increased c-Fos expression in the dBNST of DBA/1 mice. Furthermore, disruption of synaptic output from the dBNST via viral-induced tetanus neurotoxin significantly improved survival following S-IRA in DBA/1 mice without affecting baseline breathing or hypercapnic and hypoxic ventilatory response. This disruption in the dBNST resulted in changes to the balance of excitatory/inhibitory synaptic events in the downstream brainstem regions of the lateral parabrachial nucleus (PBN) and the periaqueductal gray (PAG). These findings suggest that the dBNST is a potential subcortical forebrain site necessary for the mediation of seizure-induced respiratory arrest, potentially through its outputs to brainstem respiratory regions.

**SUMMARY STATEMENT:** This study used a viral expression technique to disrupt synaptic output in the bed nucleus of the stria terminalis (BNST) of DBA/1 audiogenic seizure mice. Inactivating the BNST significantly improved survival following seizures and altered brainstem excitation/inhibition balance.

## INTRODUCTION

Sudden unexpected death in epilepsy (SUDEP) is the leading cause of death in refractory epilepsy patients (Sveinsson *et al*. 2017). Due to its unpredictable and largely unwitnessed nature, SUDEP is poorly understood and its pathophysiological mechanisms remain unknown. SUDEP cases captured in monitored settings reveal that onset of apnea occurs prior to cardiac dysfunction and death (Ryvlin *et al*. 2013), with SUDEP in rodent models following this same sequence of events (Jefferys *et al*. 2019; Lertwittayanon *et al*. 2020). This suggests that seizure-related respiratory dysfunction may play an important role in the mechanism of SUDEP.

Recent human studies have shown that stimulation of the amygdala can induce apneas, and onset of apnea is closely related to seizure spread to the amygdala (Nobis *et al*. 2019; Nobis *et al*. 2018; Dlouhy *et al*. 2015; Lacuey *et al*. 2017; Rhone *et al*. 2020). In particular, stimulation of certain portions of amygdala subnuclei may be more likely to produce apneas, with the lateral and basal areas not involved in apnea production (Nobis *et al*. 2019; Rhone *et al*. 2020). This implicates amygdalar regions associated with the extended amygdala complex, which is made up in part of the bed nucleus of the stria terminalis (BNST) and the medial and central amygdala (CeA). Because extended amygdala regions have dense reciprocal connections to brainstem respiratory, autonomic, and arousal-related nuclei, these areas are strong candidates to be involved in the neural circuit underlying seizure-related apneas.

Prior studies have shown that lesioning of the CeA in mouse models of SUDEP, including DBA/1 and Dravet Syndrome mice, can increase survival following seizures and reduce the risk of apnea during seizures (Bravo *et al*. 2020; Marincovich *et al*. 2019). The BNST is highly interconnected to the CeA (Partridge *et al*. 2016; Gungor *et al*. 2015; Pedersen *et al*. 2020) and has dense projections to respiratory-related nuclei, including the parabrachial nucleus (PBN), periaqueductal gray (PAG), ventral-lateral portion of the medulla (VLM), nucleus tractus solitarius (NTS), and serotonergic neurons in the dorsal and midbrain raphe nuclei (Dong and Swanson 2006; Dong and Swanson 2004; Pollak Dorocic *et al*. 2014). This suggests that the BNST may be another critical region involved in respiratory dysfunction during seizures.

Importantly, optogenetic stimulation of the dorsal BNST (dBNST) decreases respiratory rate (Kim *et al*. 2013), further supporting that this subcortical region may play an important role in modulating respiration. dBNST neurons, especially those projecting to the respiratory-related parabrachial nucleus (PBN), also express changes in excitability in a Dravet Syndrome mouse model, which has an increased incidence of sudden death (Yan *et al*. 2021). Altogether, these findings suggest that the dBNST may play an important role in seizure-related respiratory dysfunction and death.

In this study, we aimed to determine whether the dBNST is involved in seizure-induced respiratory arrest (S-IRA) in the DBA/1 mouse model of SUDEP. DBA/1 mice have audiogenic seizures (AGS) when exposed to a loud broad tone, and these seizures are followed by a very high incidence of S-IRA, cardiac arrest, and sudden death (Marincovich *et al*. 2019; Zhang *et al*.2018; Kommajosyula and Faingold 2019; Kommajosyula *et al*. 2017; Faingold *et al*. 2010; Schilling *et al*. 2019), making DBA/1 mice a reliable model of SUDEP.

To study the role of the dBNST in S-IRA, we first analyzed neuronal activation of the dBNST following S-IRA through immunohistochemical methods. We then used a viral-mediated strategy to disrupt synaptic transmission in the dBNST to promote survival following seizures in these mice. Finally, we examined whether lesioning of the dBNST affected breathing, chemoreception, or excitatory-inhibitory balance in brainstem regions.

## METHODS

### Mouse husbandry and genotyping

All experiments conducted with live mice were reviewed and approved by the Vanderbilt Institutional Animal Care and Use Committee, approval number M1800205-00. Animal care and experimental procedures were carried out in accordance with the National Institutes of Health Guide for the Care and Use of Laboratory Animals. DBA/1 mice were obtained from The Jackson Laboratory (Bar Harbor, ME) and Envigo (Indianapolis, IN) and group-housed under standard laboratory conditions (14 h light/10 h dark) in the mouse facility. Mice were provided food and water ad libitum.

### Priming for audiogenic seizures

DBA/1 mice were primed to have audiogenic seizures. Mice were exposed to a broad tone at 100 dB using an alarm bell (Floyd Bell, Columbus, OH) once daily for 1 min. Priming was continued until mice had full tonic-clonic seizures with respiratory arrest for a minimum of three days or until mice were primed for a full week. Following respiratory arrest, mice were resuscitated using a manual ventilator constructed in the lab. Fully primed adult mice were used in experiments and retested 3 weeks later to evaluate survival following S-IRA. Mice that did not have 3 seizures were not included in the survival study.

### c-Fos fluorescent immunohistochemistry

After DBA/1 mice were primed for a week, mice that did not exhibit 3 seizures were exposed to the audio tone again the next day, and the response was recorded as a tonic-clonic seizure with respiratory arrest, wild-running seizure, or no response. Mice were then transcardially perfused 90 minutes after audio exposure. Perfusion and immunohistochemistry were conducted as previously described to stain for c-Fos (Yan *et al*. 2021).

### Stereotaxic surgeries

Mice were anesthetized using isoflurane (initial dose: 3%; maintenance dose: 1.5%) and injected intracranially with AAV-DJ-CMV-eGFP-2A-TeNT or AAV-DJ-CMV-hrGFP (Stanford Neuroscience Gene Vector and Virus Core, Palo Alto, CA). Targeted microinjections were made bilaterally into the dBNST (coordinates from Bregma, medial/lateral: ±1.0 mm, anterior/posterior: 0.14 mm, dorsal/ventral: −4.2 mm). Mice were treated with Meloxicam for 48 h following surgery.

### Whole-body plethysmography

Whole-body plethysmography chambers (DSI, Harvard Biosciences, Inc.) were used to capture breathing changes during S-IRA in bilaterally injected mice in a subset of animals. Three weeks after stereotaxic injections, mice were placed into a plethysmography chamber and exposed to a loud broad tone to induce audiogenic seizures. Traces were plotted in Graphpad Prism.

Whole-body plethysmography was also used to evaluate changes in baseline breathing and in hypoxic and hypercapnic conditions three weeks post-injection. On the first day of trials, mice were acclimated in the plethysmography chambers for 50 min. Approximately 24 h later, baseline breathing was recorded for 50 min. On the third day, baseline breathing was recorded for 30 min, then hypercapnia (5% CO_2_) was induced for 8 min followed by a 5 min washout period. On the final day, following a 30 min baseline recording, hypoxia (8% O_2_) was induced for 8 min with a 5 min washout period after. Respiratory data (minute ventilation, breaths per minute, tidal volume) was analyzed using the DSI software and a custom-written Python script.

### Verification of injection location

Animals that survived the audiogenic-induced seizure were deeply anesthetized and transcardially perfused with ice-cold PBS followed by 4% PFA. For animals that did not survive the seizure or were used for electrophysiology, the brain was removed and fixed in 4% PFA.

Brains were coronally sectioned (30 μm) using a vibratome (Leica Biosytems, Wetzlar, Germany) and slices containing the BNST were mounted on microscope slides. Slides were then visualized on a Leica DFC290 microscope (Leica Biosystems, Wetzlar, Germany) to verify the viral expression. SigmaPlot was used to generate a heat map showing viral expression of the AAV-TeNT virus in bilaterally injected mice that survived.

### Acute slice preparation

Acute brain slices were prepared from DBA/1 mice 3-18 weeks following stereotaxic injection of AAV-DJ-CMV-eGFP-2A-TeNT or AAV-DJ-CMV-hrGFP. Animals of both sex were used and in accordance with Institutional Animal Care and Use Committee-approved protocols. Slices were prepared as previously described (Yan *et al*. 2021). Briefly, mice were anesthetized under isoflurane and decapitated. The brain was removed and coronal slices (200 μm) containing either the parabrachial nucleus (PBN) or the periaqueductal gray (PAG) were sectioned on a vibratome (Leica VT1200S, Wetzlar, Germany) in ice-cold oxygenated sucrose solution (in mM): 85 NaCl, 2.5 KCl, 1.25 NaH_2_PO_4_, 25 NaHCO_3_, 25 glucose, 75 sucrose, 0.5 sodium ascorbate, 0.1 kynurenic acid, 0.01 DL-APV, 0.5 CaCl_2_, 4.0 MgCl_2_, pH 7.39-7.42, bubbled with 95% O_2_/5% CO_2_. Slices were kept in oxygenated sucrose solution in a holding chamber at 37°C and bubbled with 95% O_2_/5% CO_2_ for at least 30 minutes. Slices were then transferred to a holding chamber containing oxygenated artificial cerebrospinal fluid (ACSF) at room temperature containing (in mM) 125 NaCl, 2.4 KCl, 1.2 NaH_2_PO_4_, NaHCO_3_, 25 glucose, pH 7.4 bubbled with 95% O_2_/5% CO_2_ and maintained under these conditions until transferred to a submerged recording chamber for recording.

### Electrophysiological recordings

Slices were perfused with oxygenated ACSF at 2.0 ml/min and maintained at 34°C. Lateral PAG (lPAG) and lateral PBN (lPBN) neurons were identified using differential interference contrast on a Slicescope II microscope (Scientifica) and whole-cell patch-clamp recordings performed using borosilicate glass micropipettes (2-8 MΩ). Signals were acquired using an Axon Multiclamp 700B amplifier (Molecular Devices) and the data was sampled at 10 kHz with a gain of 1 and a Bessel low-pass filter of 3 or 10 kHz. Series resistance was uncompensated. Access resistances were continuously monitored at 3-18 MΩ and changes >20% from the initial value were excluded from data analyses. Data were collected and analyzed using Clampex 11.1 (Molecular Devices).

Voltage clamp was performed using a cesium-methanesulfonate intracellular solution containing (in mM) 135 Cs-methanesulfonate, 10 KCl, 1 MgCl_2_, 20 sodium-phosphocreatine, 5.0 QX-314, 0.2 EGTA, 4.0 Mg-ATP, 0.3 Na-ATP, pH. 7.3, 290 mOsm. Upon break-in, cells were held at a membrane potential of −70 mV for 2 min to check for cell health and seal stability before recording the access resistance. For spontaneous excitatory postsynaptic currents (sEPSCs), cells were held at a membrane potential of −70 mV and signals recorded for 1-2 min. Following this, cells were slowly ramped to a membrane potential of +10 mV and spontaneous inhibitory postsynaptic currents (sIPSCs) recorded for 1-2 min. At the end of the recording, cells were ramped to a membrane potential of −70 mV and access resistance re-measured. Cells that experienced a change in access resistance of more than 20%, had fewer than 100 events during recording, or were unstable before both sEPSCs and sIPSCS were recorded, were disqualified from analysis.

### Experimental design and statistical analysis

Number of animals used and number of cells evaluated for each experiment are included in the Results section. Animals of either sex were used, and potential sex-specific differences were evaluated. Mean±SEM values are included throughout the text. Graphical data is depicted with mean and SEM. Outliers are noted in the text and were removed if values were outside the 95% confidence interval.

To analyze survival rates after re-induction of AGS, only animals injected with the AAV-DJ-CMV-eGFP-2A-TeNT virus with accurate bilateral targeting of the dBNST and good expression of the virus were included; animals with unilateral targeting, poor/no expression, or inaccurate targeting of the dBNST were excluded from analysis in this section. In the whole-body plethysmography and electrophysiology analyses, animals that received AAV-DJ-CMV-eGFP-2A-TeNT but whose injections either missed the dBNST, were unilateral, or had poor expression were used as part of the control group along with the animals injected with the AAV-DJ-CMV-hrGFP virus.

For the electrophysiology data, sEPSCs and sIPSCs were analyzed using MiniAnalysis (Synaptosoft). Signals at least twice the root-mean square of the noise were analyzed for amplitude and inter-event interval. Cumulative probability curves of sEPSC and sIPSC amplitudes and inter-event intervals (IEIs) were generated from the data and compared using the Kolmogorov-Smirnov (K-S) test. Mean sEPSC and sIPSC amplitude and frequency were calculated and compared using Mann-Whitney test. A p-value of <0.05 was considered statistically significant.

## RESULTS

### c-Fos expression in the dBNST significantly increases following S-IRA

To investigate how seizure-induced respiratory arrest (S-IRA) affects dBNST neuronal activation, we induced audiogenic seizures in DBA/1 mice and quantified expression of the immediate early gene c-Fos (Fig. 1A). Mice had one of three responses following audiogenic priming: no response (NR), wild-running (WR), or tonic-clonic (TC) seizure with S-IRA (followed by resuscitation). Immunohistochemistry staining revealed that animals responding with TC seizures had significantly more c-Fos positive cells in the dBNST compared to animals responding with WR seizures or NR (NR: 398.5±42.3 cells/mm^2^, n=8; WR: 387.9±33.1 cells/mm^2^, n=9; TC: 864.5±70.8 cells/mm^2^, n=14; NR vs TC: p=0.0001; WR vs TC: p<0.0001; Fig. 1B,C). We found no difference in c-Fos expression between animals with WR and NR (NR vs WR: p=0.84; Fig. 1B,C), suggesting that the significant increase in activation may result from a seizure followed by respiratory arrest.

**Figure 1:**
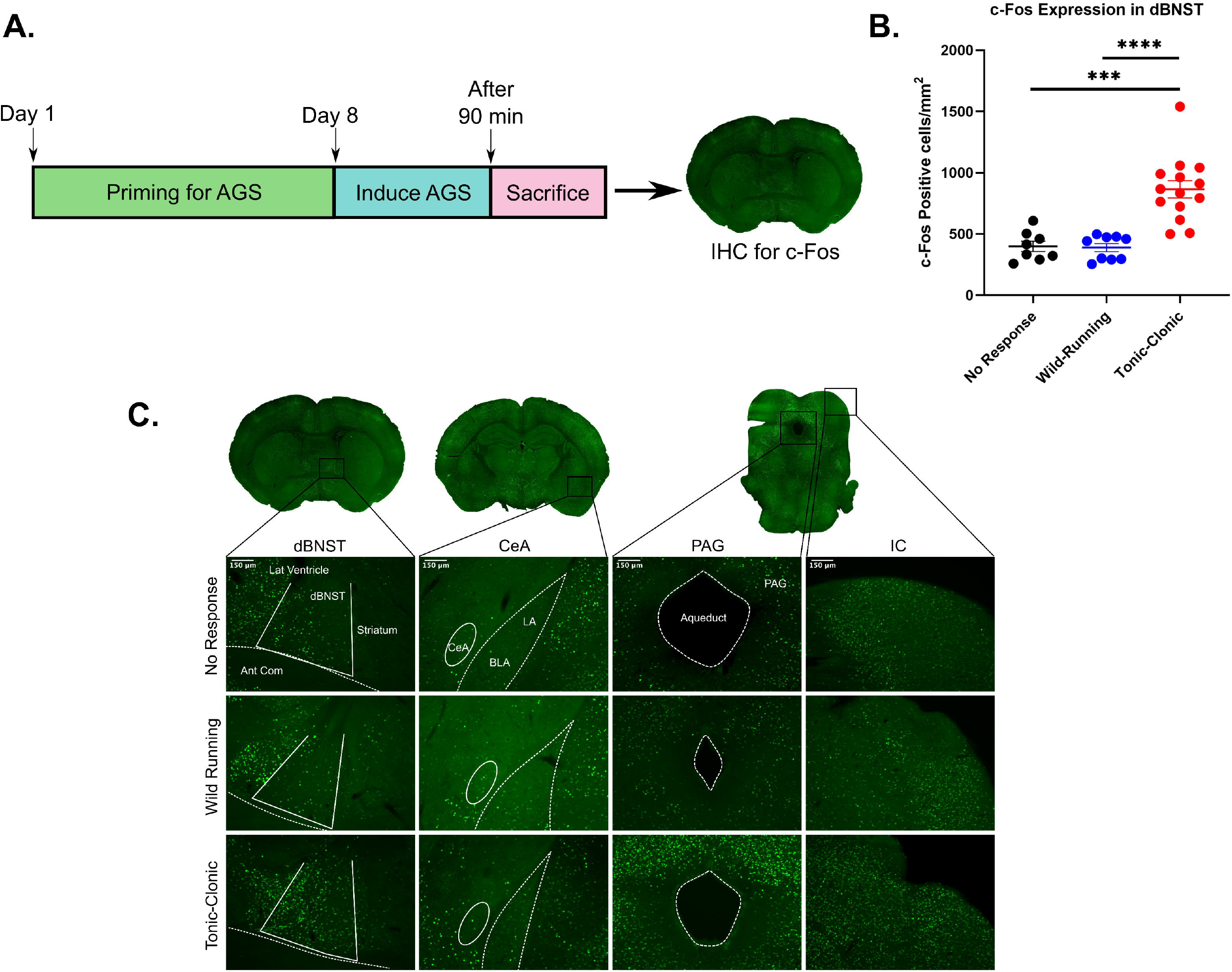
dBNST c-Fos expression significantly increases following seizure-induced respiratory arrest. **A.** Schematic showing timeline for priming DBA/1 mice for seizures. On day 8, animals were primed resulting in no response (NR), wild running (WR), or tonic-clonic seizure with S-IRA (TC), followed by immunohistochemistry for c-Fos in dBNST of mice. **B.** Scatter plot depicting significant differences in c-Fos positive cells/mm^2^ in dBNST sections for NR vs TC and WR vs TC (NR: n=8; WR: n=9; TC: n=14; NR vs TC: p=0.0001, WR vs TC: p<0.0001). No difference in c-Fos expression between NR and WR (p=0.84). **C.** Coronal sections of dBNST, CeA, PAG, and IC regions. Representative images of c-Fos expression in mice with NR, WR, and TC response 1.5 h post-priming. ***p<0.001, ****p<0.0001

In addition to the dBNST, the CeA was stained for c-Fos expression since previous studies show that electrolytic lesioning of this area can significantly increase survival after S-IRA (Bravo *et al*. 2020; Marincovich *et al*. 2019). However, we found little expression in all conditions (Fig. 1C). We also looked at c-Fos expression in the periaqueductal gray (PAG), which has been shown to have increased activity after S-IRA in DBA/1 mice (Kommajosyula *et al*. 2017), and we found greater expression in mice with S-IRA, consistent with reports in the literature (Fig. 1C). Finally, the inferior colliculus was stained as a positive control since this brain region is part of the auditory pathway and, as previously reported, shows consistent c-Fos expression with all seizure phenotypes of audiogenic response (Fig. 1C). Overall, the significant increase in dBNST c-Fos expression after S-IRA suggests that this region of the extended amygdala is likely involved in S-IRA.

### Disruption of presynaptic transmission in dBNST neurons reduces death from S-IRA in DBA/1 mice

With evidence to suggest that the dBNST may be activated in S-IRA, we next aimed to examine how disruption of presynaptic transmission in dBNST neurons would affect incidence of S-IRA and death in DBA/1 mice. We first primed the mice to have audiogenic seizures (AGS) by exposing them to a broad tone until being fully primed or for up to 7 consecutive days (Fig. 2A). Animals were considered fully primed after having 3 convulsive seizures that required resuscitation via mechanical ventilation. We then bilaterally injected tetanus neurotoxin (AAV-DJ-CMV-eGFP-2A-TeNT) or a GFP control virus (AAV-DJ-CMV-hrGFP) into the dBNST of fully primed DBA/1 mice (Fig. 2B). Electrophysiology was used to confirm that viral-induced expression of tetanus neurotoxin (AAV-TeNT) successfully resulted in disruption of presynaptic transmission in dBNST neurons compared to the GFP control virus (AAV-GFP). sIPSC frequency was significantly reduced in cells expressing the AAV-TeNT virus compared to the AAV-GFP virus (AAV-GFP: 4.0±1.1 Hz, n=5 cells from 3 animals; AAV-TeNT: 1.0±0.33 Hz, n=7 cells from 3 animals; p=0.01; Fig. 2D). There was no significant change in sIPSC amplitude in the AAV-TeNT expressing cells (AAV-GFP: 23.6±4.0 pA, n=5 cells from 3 animals; AAV-TeNT: 22.2±2.8 pA, n=7 cells from 3 animals; p=0.76; Fig. 2E).

**Figure 2:**
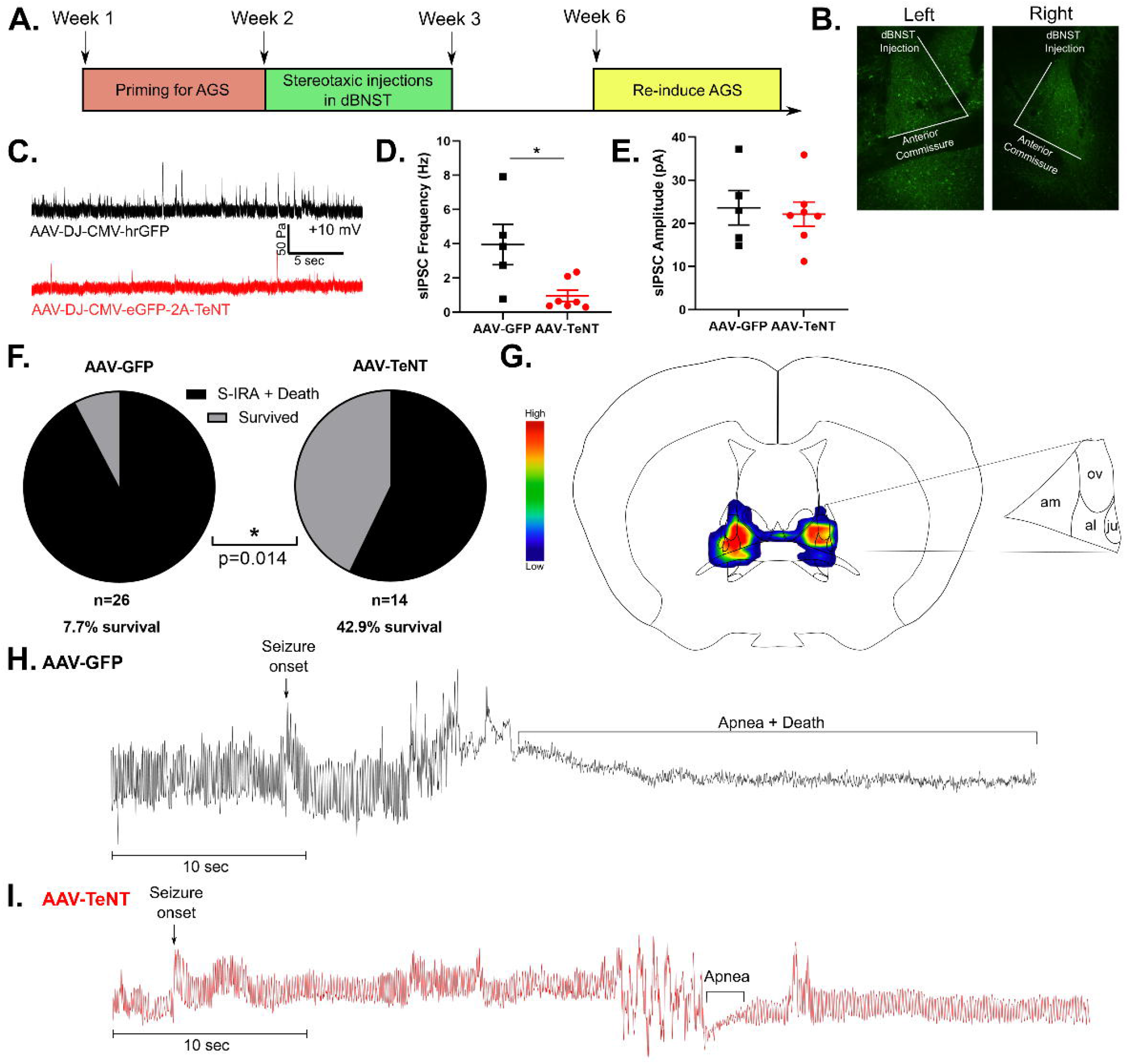
Disrupting presynaptic transmission in dBNST neurons reduces seizure-induced death in DBA/1 mice. **A.** Schematic showing timeline for priming DBA/1 mice for seizures, followed by stereotaxic injections and re-induction of seizures to observe survival rate. **B.** Representative image showing bilateral viral expression of AAV-DJ-CMV-eGFP-2A-TeNT in left and right dBNST. **C.** Representative trace of sIPSCs in dBNST neurons of animals with viral expression of AAV-GFP vs. AAV-TeNT. **D.** Graph shows significant reduction in sIPSC frequency in cells expressing AAV-TeNT (AAV-GFP: n=5 cells from 3 animals; AAV-TeNT: n=7 cells from 3 animals; p=0.01). **E.** Graph showing no difference in sIPSC amplitude between AAV-GFP and AAV-TeNT expressing cells (AAV-GFP: n=5 cells from 3 animals; AAV-TeNT: n=7 cells from 3 animals; p=0.76). **F.** Graph depicting significant increase in survival following SIRA in AAV-TeNT virus-injected animals compared to AAV-GFP animals (AAV-GFP: n=26; AAV-TeNT: n=14; p=0.014). **G.** Heat map on atlas image shows AAV-TeNT viral expression in dBNST for mice that survived (am: anteromedial area, ov: oval nucleus, al: anterolateral area, ju: juxtacapsular nucleus). **H.** Representative plethysmography trace in AAV-GFP animal showing S-IRA followed by death. **I.** Representative plethysmography trace in AAV-TeNT animal depicting survival following S-IRA. *p<0.05

Three weeks following stereotaxic injection in the dBNST, we re-induced seizures in these mice to observe if there were any differences in incidence of S-IRA and survival between the AAV-TeNT and AAV-GFP control mice. We found a significant increase in survival following disruption of presynaptic transmission in the dBNST. 42.9% of AAV-TeNT virally injected mice survived while only 7.7% of AAV-GFP mice survived (AAV-GFP: n=26; AAV-TeNT: n=14; p=0.014; Fig. 2F). Unilateral targeting of the dBNST was insufficient to prevent death in AAV-TeNT mice, and only 4.3% of animals with unilateral targeting survived (n=23), similar to the survival rate in the AAV-GFP group. Thus, the AAV-TeNT group only consisted of animals with accurate bilateral targeting of the dBNST. In mice that survived, bilateral viral expression of AAV-TeNT was greatest in the anteromedial area of the dBNST, followed by high levels of expression in the oval nucleus and anterolateral area (Fig. 2G).

Using whole-body plethysmography to visualize breathing during S-IRA, we found that in AAV-GFP mice that died from S-IRA, seizures were followed by apnea and death (Fig. 2H). In contrast, AAV-TeNT mice that survived experienced seizures that resulted in short apneas followed by return to normal breathing (Fig. 2I).

Animals that did not have seizures three weeks following the stereotaxic injection were excluded from survival analysis. There was no relationship to the type of viral injection in those animals that were non-responders to audio tone. These animals may have lost seizure susceptibility due to placement of ear bars during surgery, damaging their eardrums and decreasing their ability to perceive sound and thus have an audiogenic seizure. Susceptibility to audiogenic seizures changes and often diminishes as DBA/1 mice age, so age could also explain why we could not re-induce seizures in some fully primed mice (Martin *et al*. 2020).

### Disruption of synaptic output in the dBNST does not alter breathing or chemoreception

To determine if disrupting synaptic output of the dBNST affects breathing or chemoreception, we used whole-body plethysmography to analyze baseline breathing, hypercapnic ventilatory response (HCVR), and hypoxic ventilatory response (HVR). Animals with the AAV-GFP virus and poor targeting or expression of the AAV-TeNT virus were considered as part of the control group. There were no changes in baseline breathing respiratory rate (Control: 100±5.1%, n=29; TeNT: 105.7±9.2%, n=16; p=0.56; Fig. 3A), minute ventilation (Control: 100±5.9%, n=29; TeNT: 100.7±8.4%, n=16; p=0.95; Fig. 3B), or tidal volume (Control: 100±6.3%, n=29; TeNT: 95.4±8.5%, n=16; p=0.66; Fig. 3C) between control and AAV-TeNT mice.

**Figure 3:**
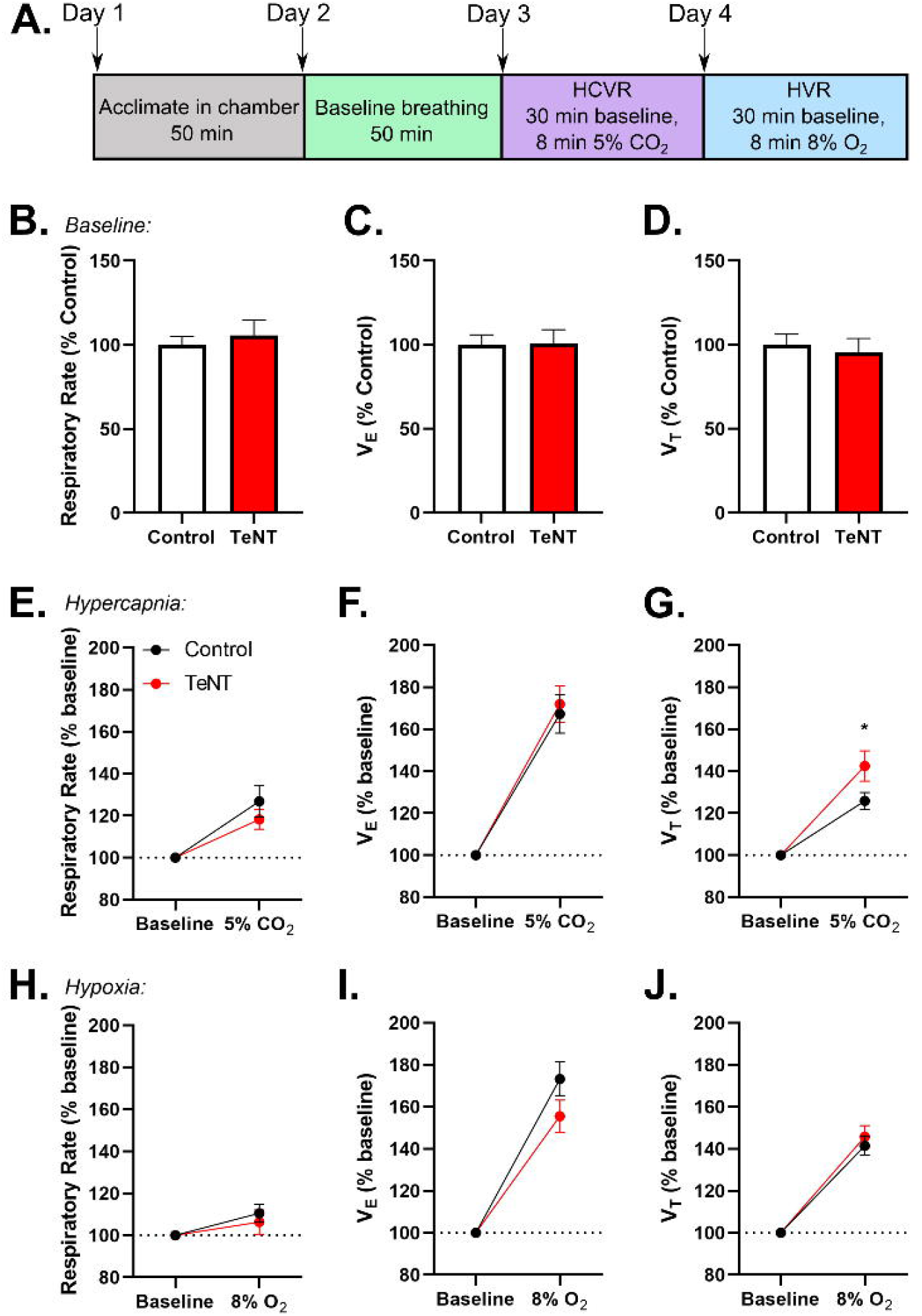
Disrupting synaptic output in dBNST neurons does not significantly alter breathing or chemoreception. **A.** Schematic showing experimental design for whole-body plethysmography trials. No difference in baseline breathing **B.** respiratory rate (p=0.56), **C.** V_E_ (p=0.95), or **D.** V_T_ (p=0.66) between control and AAV-TeNT injected animals. No difference in **E.** respiratory rate (p=0.41) and **F.** V_E_ (p=0.73). AAV-TeNT animals had a significant increase in **G.** V_T_ (p=0.035) during hypercapnic conditions. No changes in HVR **H.** respiratory rate (p=0.56), **I.** V_E_ (p=0.16), or **J.** V_T_ (p=0.56). *p<0.05 (V_E_: minute ventilation; V_T_: tidal volume)

We found no differences between control and AAV-TeNT animals in HCVR respiratory rate (Control: 126.8±7.5%, n=27 with 2 high outliers removed; TeNT: 118.2±4.7%, n=16; p=0.41; Fig. 3D) and minute ventilation (Control: 167.2±9.1%, n=29; TeNT: 172±8.6%, n=16; p=0.73; Fig. 3E), but there was a significant increase in tidal volume (Control: 125.8±4.1%, n=29; TeNT: 142.4±7.2%, n=16; p=0.035; Fig. 3F) of AAV-TeNT animals. Increased CO_2_ can induce panic behavior in mice, and this response is dependent on BNST function (Taugher *et al*. 2014). Therefore, the increase in tidal volume in the AAV-TeNT animals may represent blunted panic behavior due to hypofunctioning of the BNST, with an otherwise preserved chemosensory response.

Finally, there were no significant differences in HVR respiratory rate (Control: 110.6±4.4%, n=26 with 3 high outliers removed; TeNT: 106.3±5.9% n=16; p=0.56; Fig. 3G), minute ventilation (Control: 173.3±8.1%, n=26; TeNT: 155.5±7.8%, n=16; p=0.16; Fig. 3H), or tidal volume (Control: 141.5%±4.4%, n=29; TeNT: 145.7±5.4%, n=16; p=0.56; Fig. 3I) between control and AAV-TeNT mice. Overall, these results suggest that the dBNST is not involved in baseline breathing or respiration in response to chemoreception.

### Disrupting presynaptic transmission from the dBNST neurons altered excitatory synaptic drive in the downstream brainstem region of the lateral parabrachial nucleus

Given the disruption in S-IRA and the BNST projections to brainstem respiratory regions, we wanted to determine if our manipulation had any effect on these end target regions of interest. We chose to investigate the lPBN and the lPAG as they both receive dense input from the extended amygdala, are involved in respiratory control, and have been shown to be activated by S-IRA induced death in DBA/1 mice (Kommajosyula *et al*. 2017; Faull *et al*. 2019; Holstege *et al*. 1985; Luskin *et al*. 2021; Dutschmann and Herbert 2006). To elucidate if dBNST inactivation altered E/I balance in these brainstem neurons, we recorded sEPSCs and sIPSCs in both the lPBN and lPAG neurons from control (n=6 animals for both lPBN and lPAG recordings) and AAV-TeNT (n=3 and 5 animals for lPBN and lPAG recordings, respectively) animals.

Based on the recordings in the lPBN, we observed a reduction in mean sEPSC amplitude in AAV-TeNT animals compared to controls (Control: 13.3±1.2 pA, n=6 cells; TeNT: 10.4±0.3 pA, n=5 cells; p=.03; Fig. 4B) with a leftward shift in the distribution of sEPSC amplitudes from AAV-TeNT animals compared to controls (K-S: p<0.0001; Fig. 4F), denoting a reduction in the size of synaptic excitatory events on lPBN neurons. This reduction in amplitude was observed without changes in mean sEPSC frequency (Control: 7.8±1.5 Hz; TeNT: 6.2±0.89 Hz, n=5; p=0.54; Fig. 4C), though we did observe a significant rightward shift in the distribution of IEIs (K-S: p<0.0001; Fig. 4G), overall indicating a significant reduction in excitatory synaptic drive in the lPBN.

**Figure 4:**
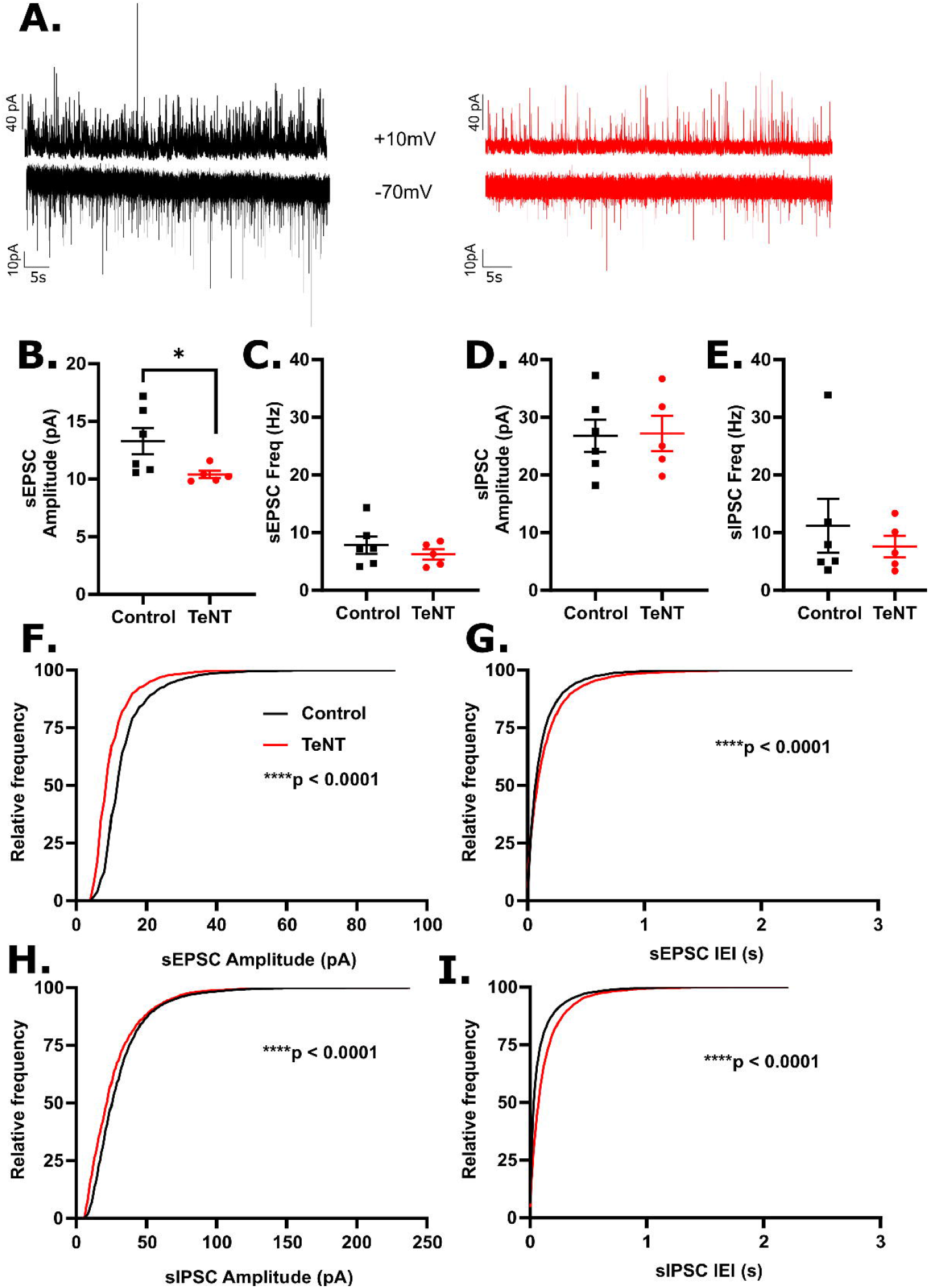
Disrupting synaptic output in dBNST neurons reduced excitatory synaptic drive at lPBN neurons. **A.** Examples of recordings of sEPSCs (bottom) and sIPSCs (top) recorded from control (black) and AAV-TeNT (red) animals (n=6 and 3 animals, respectively). Each trace is 60s in duration. **B.** Mean sEPSC amplitude was significantly reduced in neurons from AAV-TeNT animals compared to control animals (control: n=6 cells; TeNT: n=5 cells; p=.03), though no effect was observed on **C.** mean frequency (p=0.54). No effect of dBNST synaptic disruption was observed on **D.** mean sIPSC amplitude (p=0.93) or **E.** mean frequency (p=0.79) **F.** A significant leftward shift in the distribution of sEPSC amplitudes (K-S: p<0.0001) was observed as was **G.** a significant rightward shift in the distribution of sEPSC frequencies (K-S: p<0.0001). **H.** A significant leftward shift in the distribution of sIPSC amplitudes (K:S: p<0.0001) and **I.** a significant rightward shift in the distribution sIPSC frequencies (K-S: p<0.0001) were observed. *p<0.05, ****p<0.0001

Though we did not observe any differences in mean sIPSC amplitude (Control: 26.7±2.8 pA, n=6 cells; TeNT: 27.2±3.1 pA, n=5 cells; p=0.93; Fig. 4D) or frequency (Control: 11.2±4.7 Hz, n=6 cells; TeNT: 7.6±1.8 Hz, n=5 cells; p=0.79; Fig. 4E), we did observe similar changes in the distributions of sIPSC amplitudes and IEIs as we did with the sEPSCs: a leftward shift in amplitudes (K-S: p<0.0001; Fig. 4H) and a rightward shift in IEIs (K-S: p<0.0001; Fig. 4I) suggesting reductions in populations of inhibitory signaling in the lPBN following our manipulation.

Unlike the lPBN, the synaptic changes were more subtle in the lPAG following AAV-TeNT injections in the dBNST. Most noticeably, we observed a significant rightward shift in the distribution of sIPSC amplitudes (K-S: p<0.0001; Fig. 5H) but no change in the mean sIPSC amplitudes (Control: 28.9±2.6 pA, n=9 cells from 6 animals; TeNT: 33.4±10.6 pA, n=6 cells from 5 animals; p=1.0; Fig. 5D), suggesting minor changes in the distribution of synaptic inhibitory size that occurred with no change in mean frequency of events (Control: 3.3±0.7 Hz; TeNT: 3.3±0.7 Hz; p=0.86; Fig. 5E) with a subtle leftward shift on cumulative probability plot (K-S: p<0.01, Fig. 5I).

**Figure 5:**
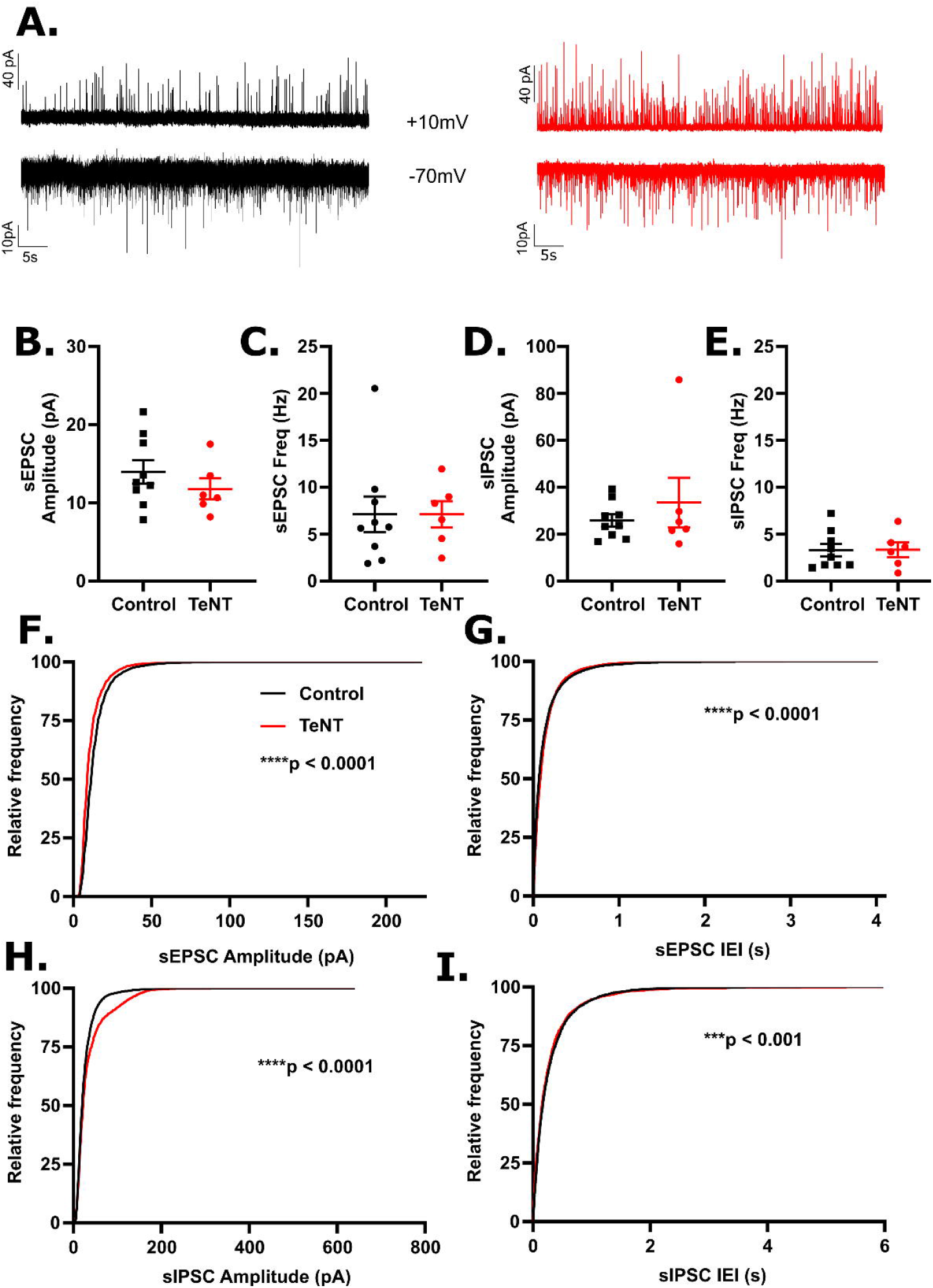
Disrupting dBNST synaptic output to lPAG neurons resulted in subtle changes to excitatory and inhibitory signaling. **A.** Examples of recordings of sEPSCs (bottom) and sIPSCs (top) recorded from control (black) and AAV-TeNT (red) animals (n=6 and 5 animals, respectively). Each trace is 60s in duration. No effect of dBNST synaptic disruption was observed on **B.** mean sEPSC amplitude (Control: n=9 cells; TeNT: n=6 cells; p=0.33), **C.** mean sEPSC frequency (p=0.61), **D.** mean sIPSC amplitude (p=1.0), or **E.** mean sIPSC frequency (p=0.86). **F.** A leftward shift in the distribution of sEPSC amplitudes (K-S: p<0.0001) and G. a rightward shift in the distribution of sIPSC frequencies (K-S: p<0.0001) were observed. **H.** A significant rightward shift in the distribution of sIPSC amplitudes (K-S: p<0.0001) and **I.** a significant leftward shift in the distribution of sIPSC frequencies (K-S: p<0.0001) were observed. ****p<0.0001

Likewise, there were no differences in mean sEPSC amplitude (Control: 14.0±1.5 pA; TeNT: 11.8±1.3 pA, n=6 cells from 5 animals; p=0.33; Fig. 5B) or frequency (Control: 7.1±1.9 Hz, n=9 cells from 6 animals; TeNT: 7.1±1.4 Hz, n=6 cells from 5 animals; p=0.61; Fig. 5C) and only very slight, but statistically significant, shifts in the distribution of sEPSC amplitudes (leftward shift; K-S: p<0.0001; Fig. 5F) and sEPSC IEIs (rightward shift; K-S: p<0.0001; Fig. 5G).

Together, our data shows that this chronic synaptic disruption of the dBNST can have effects in the downstream brainstem target regions of the lPBN and the lPAG. In general, these effects were more pronounced in the lPBN where we observed significantly reduced sEPSC amplitudes, suggesting that disruption of dBNST input is producing postsynaptic changes in these neurons. The dBNST has both excitatory and inhibitory projections to the lPBN (Luskin *et al*. 2021), so these reductions in sEPSC amplitudes could be due to a loss of the excitatory projection and/or compensatory change for loss of the inhibitory input. The heterogeneous nature of BNST neurons, the relative variability in targeting, and the time for compensatory changes to develop likely contribute to the overall subtle effects.

## DISCUSSION

Here we demonstrated that bilateral disruption of synaptic transmission in the dBNST can greatly reduce S-IRA and death in the DBA/1 audiogenic seizure mouse model of SUDEP. Prior work elegantly demonstrated through lesion studies that the amygdala may be necessary for seizure-induced respiratory arrest in DBA/1 mice. In this study, we demonstrated that the dBNST was the extended amygdalar component that was strongly activated during seizures with respiratory arrest in DBA/1 mice, as measured by expression of the immediate early gene c-Fos. By disrupting synaptic transmission in dBNST neurons with a viral-mediated approach, we were able to disrupt S-IRA and death in DBA/1 mice, suggesting that the dBNST is also a critical mediator of this effect. Breathing under normoxic, hypercapnic, and hypoxic conditions was not affected by these dBNST manipulations. Furthermore, we investigated the influence that the relative dBNST silencing had on E/I balance in downstream brainstem respiratory regions, the lPBN and lPAG. Here we observed that we could significantly alter synaptic signaling in these regions; in particular, the lPBN was more greatly affected with evidence of reduction in excitatory signaling in this critical target region.

There is building evidence that seizure spread to, or electrical stimulation of, the amygdala can induce apneas and respiratory dysfunction in patients with epilepsy (Nobis *et al*. 2019; Nobis *et al*. 2018; Dlouhy *et al*. 2015; Rhone *et al*. 2020). Central apneas during seizure onset have unclear clinical impact (Tio *et al*. 2020), with respiratory dysfunction and apneas persisting after seizures being potentially more clinically impactful (Vilella *et al*. 2018). The S-IRA present in the DBA/1 mouse model consists of a rapid-onset apnea during the clonic phase of the seizure that persists with ultimate hypoventilation and death (Schilling *et al*. 2019). This represents a potential model for both ictal apnea and an extended postictal apnea.

Seizure-induced death in DBA/1 mice also activates brainstem regions (Kommajosyula *et al*.2017). Growing evidence from chemoconvulsant seizure models suggests spread of seizures into the brainstem are associated with apneas and death (Jefferys *et al*. 2019; Lertwittayanon *et al*. 2020) and that brainstem seizures can be associated with brainstem spreading depolarization (Jansen *et al*. 2019). We propose that prior clinical and preclinical evidence, along with our current study, suggest that the extended amygdala is a potential node for seizure propagation to brainstem respiratory structures, producing central apneas and also impairing respiratory rescue responses and ultimately increasing the risk for SUDEP.

This study appears to stand in contrast to the work implicating the central amygdala (CeA) as necessary for S-IRA (Marincovich *et al*. 2019). However, given the highly connected nature of the CeA and BNST, it is not unexpected that we have similar results with manipulations to the dBNST. The clinical stimulation data clearly implicates a region in the centromedial amygdala as an area that can readily produce an apnea, but it is also apparent from preclinical studies that the BNST can have a similar function in modulating respiratory rate (Kim *et al*. 2013). The BNST is also involved in arousal (Kodani *et al*. 2017) and has projections to important chemosensory structures such as the raphe (Weissbourd *et al*. 2014), so its activation during seizures may be important in both initial apnea generation but also in continued respiratory and arousal deficits with seizures. The BNST and CeA then may be working in concert in a pathological way to increase SUDEP risk if either or both are activated by a seizure.

It is unclear how often the BNST is activated by seizures as it has not been regularly evaluated in prior studies, but it’s striking that c-Fos data here mirrors previously published c-Fos data in a Dravet Syndrome mouse model that also has a significant rate of SUDEP (Yan *et al*. 2021). Seizure activation of both the CeA and BNST may be a marker of SUDEP risk but would need to be studied in more preclinical models. We did not show significant c-Fos expression in the CeA in the DBA/1 model with seizures, but this does not preclude its activation as a very small subregion may be all that is needed to induce apnea, as suggested by human data (Rhone *et al*. 2020).

Interestingly, similar to the Marincovich *et al*. 2019 study, we were not able to improve survival after S-IRA in 100% of our mice with dBNST manipulations, and we were only able to improve survival following S-IRA in mice with accurate bilateral viral targeting of the dBNST. These findings can be attributed to several factors. First, there is likely not complete viral penetration of the dBNST since the virus may not transduce into all cells of this region. The viral-mediated approach also diminishes synaptic activity rather than lesioning the dBNST, providing greater accuracy in targeted regions while causing a response that may only be exhibited with bilateral inactivation. Additionally, the BNST is a highly heterogeneous brain region; however, our approach targets the whole dBNST rather than particular subregions or projection neurons, creating variability in survival due to differences in viral expression between animals. Moreover, since the BNST and CeA are highly interconnected, both amygdalar regions may be involved in S-IRA, and targeting cells in the BNST interconnected with the CeA may be necessary for improving survival, which could explain why there is not complete survival in both this study and the Marincovich *et al*. 2019 study. Hence, a multifactorial pathology likely underlies this phenomenon and makes it challenging to prevent death by inactivating just one extended amygdala region, further implicating both the BNST and CeA as potentially important regions to target in reducing SUDEP risk.

We further began to determine which brainstem region(s) the extended amygdala is acting on to influence respiratory status. The Kölliker-Fuse nucleus of the lateral PBN and the PAG are both activated during S-IRA and also activated when S-IRA is alleviated through the actions of fluoxetine (Kommajosyula and Faingold 2019; Kommajosyula *et al*. 2017), making them likely candidate regions. We found virally-mediated inhibition of the dBNST significantly altered E/I balance in both the lPBN and the lPAG to some degree with more pronounced alterations in the lPBN, exhibiting decreased excitatory synaptic drive. Yan *et al*. 2021 previously found that BNST-PBN neurons in a mouse model of Dravet Syndrome and SUDEP were less likely to fire action potentials in response to current injections, building evidence on the potential importance of this pathway and risk for SUDEP. In Sudden Infant Death Syndrome (SIDS), another cause of sudden death due to respiratory dysfunction, and in Rett Syndrome, a neurodevelopmental disorder with significant apneas, the Köllliker-Fuse/lateral PBN have also been implicated (Abdala *et al*. 2016; Varga *et al*. 2020; Lavezzi *et al*. 2019). More broadly, stimulation of BNST-PBN projection neurons has also been shown to decrease respiratory rate (Kim et al., 2013).

Within BNST to brainstem region circuits, our BNST manipulations may also be affecting serotonergic signaling. Augmentation of serotonin (5-HT) function in DBA/1 mice promotes survival and diminishes S-IRA (Zhang *et al*. 2018; Zeng *et al*. 2015; Faingold *et al*. 2016). DBA/1 mice have defective expression of serotonergic receptors in the brainstem (Feng and Faingold 2015), which may explain the ability of 5-HT augmentation in promoting survival. However, given significant extended amygdala output to dorsal raphe neurons (Weissbourd *et al*. 2014) and the ability of activation of BNST neurons to modulate 5-HT metabolism in the DR (Donner *et al*. 2019), it is possible that the extended amygdala also has a role in 5-HT dysfunction in these animals. More recently, adrenergic α2 receptor agonists have been shown to alleviate S-IRA in DBA/1 mice (Feng and Faingold 2015; Zhang *et al*. 2021). Activation of these receptors in the BNST can modulate PBN input (Harris *et al*. 2018; Fetterly *et al*. 2018), which may provide a mechanism outside the noradrenergic role in arousal for the effect of adrenergic α2 receptor agonists.

This study further supports that a central apnea is critically important in leading to SUDEP, supporting human studies and observations (Ryvlin *et al*. 2013; Johnson *et al*. 2021). Overall, this study builds on the growing evidence for the extended amygdala as a key node that critically affects respiratory function and apnea induction but potentially has multiple levels of influence in SUDEP pathophysiology. More studies will be needed to fully test this hypothesis, in particular to begin to address the heterogeneity of this region and determine those neuronal populations that are most critically involved in S-IRA. Determination of these precise neuronal populations will aid in development of therapeutic targets to prevent death in SUDEP and other syndromic sudden death entities including SIDS, whether through neuromodulation of these extended amygdalar pathways or neuromodulation of other brain regions that may counteract the respiratory and arousal depression. More broadly, this study further defines the extended amygdala as a higher order structure that can modulate brainstem respiratory networks.

In summary, we demonstrated that a bilateral virally-mediated impairment in dBNST synaptic function greatly diminished S-IRA and altered E/I balance in brainstem respiratory regions, suggesting that the BNST may represent an important node in the pathway by which seizures spread to the brainstem to cause death in the DBA/1 audiogenic seizure model of SUDEP.

## Acknowledgements

The authors would like to acknowledge Ragan Huffman, Katie Preisinger and Cierra Lyons for assistance in experiments and Jagath Vytheeswaran for writing Python scripts for plethysmography analysis.

## REFERENCES

Abdala A. P., Toward M. A., Dutschmann M., Bissonnette J. M., Paton J. F. R. (2016) Deficiency of GABAergic synaptic inhibition in the Kölliker-Fuse area underlies respiratory dysrhythmia in a mouse model of Rett syndrome. J. Physiol. 594, 223–237.

Bravo E., Marincovich A., Teran F., Crotts M., Lefkowitz R. J., Richerson G. (2020) Postictal modulation of breathing by the central amygdala (CeA) can induce sudden unexpected death in epilepsy (SUDEP). FASEB J. 34, 1–1.

Dlouhy B. J., Gehlbach B. K., Kreple C. J., Kawasaki H., Oya H., Buzza C., Granner M. A., et al. (2015) Breathing Inhibited When Seizures Spread to the Amygdala and upon Amygdala Stimulation. J. Neurosci. 35, 10281–10289.

Dong H.-W., Swanson L. W. (2004) Organization of axonal projections from the anterolateral area of the bed nuclei of the stria terminalis. J. Comp. Neurol. 468, 277–298.

Dong H.-W., Swanson L. W. (2006) Projections from bed nuclei of the stria terminalis, anteromedial area: cerebral hemisphere integration of neuroendocrine, autonomic, and behavioral aspects of energy balance. J. Comp. Neurol. 494, 142–178.

Donner N. C., Mani S., Fitz S. D., Kienzle D. M., Shekhar A., Lowry C. A. (2019) Crh receptor priming in the bed nucleus of the stria terminalis induces tph2 gene expression in the dorsomedial dorsal raphe nucleus and chronic anxiety. Prog. Neuropsychopharmacol. Biol. Psychiatry, 109730.

Dutschmann M., Herbert H. (2006) The Kölliker-Fuse nucleus gates the postinspiratory phase of the respiratory cycle to control inspiratory off-switch and upper airway resistance in rat. Eur. J. Neurosci. 24, 1071–1084.

Faingold C. L., Randall M., Tupal S. (2010) DBA/1 mice exhibit chronic susceptibility to audiogenic seizures followed by sudden death associated with respiratory arrest. Epilepsy Behav. 17, 436–440.

Faingold C. L., Randall M., Zeng C., Peng S., Long X., Feng H.-J. (2016) Serotonergic agents act on 5-HT3 receptors in the brain to block seizure-induced respiratory arrest in the DBA/1 mouse model of SUDEP. Epilepsy Behav. 64, 166–170.

Faull O. K., Subramanian H. H., Ezra M., Pattinson K. T. S. (2019) The midbrain periaqueductal gray as an integrative and interoceptive neural structure for breathing. Neurosci. Biobehav. Rev. 98, 135–144.

Feng H.-J., Faingold C. L. (2015) Abnormalities of serotonergic neurotransmission in animal models of SUDEP. Epilepsy Behav.

Fetterly T. L., Basu A., Nabit B. P., Awad E., Williford K. M., Centanni S. W., Matthews R. T., Silberman Y., Winder D. G. (2018) α2A-adrenergic receptor activation decreases parabrachial nucleus excitatory drive onto BNST CRF neurons and reduces their activity in vivo. J. Neurosci.

Gungor N. Z., Yamamoto R., Paré D. (2015) Optogenetic study of the projections from the bed nucleus of the stria terminalis to the central amygdala. J. Neurophysiol. 114, 2903–2911.

Harris N. A., Isaac A. T., Günther A., Merkel K., Melchior J., Xu M., Eguakun E., et al. (2018) Dorsal BNST α2A-Adrenergic Receptors Produce HCN-Dependent Excitatory Actions That Initiate Anxiogenic Behaviors. J. Neurosci. 38, 8922–8942.

Holstege G., Meiners L., Tan K. (1985) Projections of the bed nucleus of the stria terminalis to the mesencephalon, pons, and medulla oblongata in the cat. Exp. Brain Res. 58, 379–391.

Jansen N. A., Schenke M., Voskuyl R. A., Thijs R. D., Maagdenberg A. M. J. M. van den, Tolner E. A. (2019) Apnea associated with brainstem seizures in Cacna1aS218L mice is caused by medullary spreading depolarization. J. Neurosci.

Jefferys J. G. R., Arafat M. A., Irazoqui P. P., Lovick T. A. (2019) Brainstem activity, apnea, and death during seizures induced by intrahippocampal kainic acid in anaesthetized rats. Epilepsia 60, 2346–2358.

Johnson M., Samudra N., Gallagher M. J., Abou-Khalil B., Nobis W. P. (2021) Near SUDEP during bilateral stereo-EEG monitoring characterized by diffuse postictal EEG suppression. Epilepsia epi.16852.

Kim S.-Y., Adhikari A., Lee S. Y., Marshel J. H., Kim C. K., Mallory C. S., Lo M., et al. (2013) Diverging neural pathways assemble a behavioural state from separable features in anxiety. Nature 496, 219–223.

Kodani S., Soya S., Sakurai T. (2017) Excitation of GABAergic Neurons in the Bed Nucleus of the Stria Terminalis Triggers Immediate Transition from Non-Rapid Eye Movement Sleep to Wakefulness in Mice. J. Neurosci. 37, 7164–7176.

Kommajosyula S. P., Faingold C. L. (2019) Neural activity in the periaqueductal gray and other specific subcortical structures is enhanced when a selective serotonin reuptake inhibitor selectively prevents seizure induced sudden death in the DBA /1 mouse model of sudden unexpected death in epilepsy.

Kommajosyula S. P., Randall M. E., Brozoski T. J., Odintsov B. M., Faingold C. L. (2017) Specific subcortical structures are activated during seizure-induced death in a model of sudden unexpected death in epilepsy (SUDEP): A manganese-enhanced magnetic resonance imaging study.

Lacuey N., Zonjy B., Londono L., Lhatoo S. D. (2017) Amygdala and hippocampus are symptomatogenic zones for central apneic seizures. Neurology 88, 701–705.

Lavezzi A. M., Ferrero S., Paradiso B., Chamitava L., Piscioli F., Pusiol T. (2019) Neuropathology of Early Sudden Infant Death Syndrome-Hypoplasia of the Pontine Kolliker-Fuse Nucleus: A Possible Marker of Unexpected Collapse during Skin-to-Skin Care. Am. J. Perinatol. 36, 460–471.

Lertwittayanon W., Devinsky O., Carlen P. L. (2020) Cardiorespiratory depression from brainstem seizure activity in freely moving rats. Neurobiol. Dis. 134, 104628.

Luskin A. T., Bhatti D. L., Mulvey B., Pedersen C. E., Girven K. S., Oden-Brunson H., Kimbell K., et al. (2021) Extended amygdala-parabrachial circuits alter threat assessment and regulate feeding. Sci Adv 7.

Marincovich A., Bravo E., Dlouhy B., Richerson G. B. (2019) Amygdala lesions reduce seizure-induced respiratory arrest in DBA/1 mice. Epilepsy Behav., 106440.

Martin B., Dieuset G., Pawluski J. L., Costet N., Biraben A. (2020) Audiogenic seizure as a model of sudden death in epilepsy: A comparative study between four inbred mouse strains from early life to adulthood. Epilepsia 61, 342–349.

Nobis W. P., González Otárula K. A., Templer J. W., Gerard E. E., VanHaerents S., Lane G., Zhou G., Rosenow. J. M., Zelano C., Schuele S. (2019) The effect of seizure spread to the amygdala on respiration and onset of ictal central apnea. J. Neurosurg. 132, 1313–1323.

Nobis W. P., Schuele S., Templer J. W., Zhou G., Lane G., Rosenow J. M., Zelano C. (2018) Amygdala-stimulation-induced apnea is attention and nasal-breathing dependent. Annals of Neurology 83, 460–471.

Partridge J. G., Forcelli P. A., Luo R., Cashdan J. M., Schulkin J., Valentino R. J., Vicini S. (2016) Stress increases GABAergic neurotransmission in CRF neurons of the central amygdala and bed nucleus stria terminalis. Neuropharmacology 107, 239–250.

Pedersen W. S., Schaefer S. M., Gresham L. K., Lee S. D., Kelly M. P., Mumford J. A., Oler J. A., Davidson R. J. (2020) Higher resting-state BNST-CeA connectivity is associated with greater corrugator supercilii reactivity to negatively valenced images. Neuroimage 207, 116428.

Pollak Dorocic I., Dorocic I. P., Fürth D., Xuan Y., Johansson Y., Pozzi L., Silberberg G., Carlén M., Meletis K. (2014) A Whole-Brain Atlas of Inputs to Serotonergic Neurons of the Dorsal and Median Raphe Nuclei.

Rhone A. E., Kovach C. K., Harmata G. I., Sullivan A. W., Tranel D., Ciliberto M. A., Howard M. A., et al. (2020) A human amygdala site that inhibits respiration and elicits apnea in pediatric epilepsy. JCI Insight 5.

Ryvlin P., Nashef L., Lhatoo S. D., Bateman L. M., Bird J., Bleasel A., Boon P., et al. (2013) Incidence and mechanisms of cardiorespiratory arrests in epilepsy monitoring units (MORTEMUS): a retrospective study. Lancet Neurol. 12, 966–977.

Schilling W. P., McGrath M. K., Yang T., Glazebrook P. A., Faingold C. L., Kunze D. L. (2019) Simultaneous cardiac and respiratory inhibition during seizure precedes death in the DBA/1 audiogenic mouse model of SUDEP. PLoS One 14, e0223468.

Sveinsson O., Andersson T., Carlsson S., Tomson T. (2017) The incidence of SUDEP: A nationwide population-based cohort study. Neurology 89, 170–177.

Taugher R. J., Lu Y., Wang Y., Kreple C. J., Ghobbeh A., Fan R., Sowers L. P., Wemmie J. A. (2014) The Bed Nucleus of the Stria Terminalis Is Critical for Anxiety-Related Behavior Evoked by CO2 and Acidosis. J. Neurosci. 34, 10247–10255.

Tio E., Culler G. W., Bachman E. M., Schuele S. (2020) Ictal central apneas in temporal lobe epilepsies. Epilepsy Behav. 112, 107434.

Varga A. G., Maletz S. N., Bateman J. T., Reid B. T., Levitt E. S. (2020) Neurochemistry of the Kölliker-Fuse nucleus from a respiratory perspective. J. Neurochem.

Vilella L., Lacuey N., Hampson J. P., Sandhya Rani M. R., Sainju R. K., Friedman D., Nei M., et al. (2018) Postconvulsive central apnea as a biomarker for sudden unexpected death in epilepsy (SUDEP). Neurology, 10.1212/WNL.0000000000006785.

Weissbourd B., Ren J., DeLoach K. E., Guenthner C. J., Miyamichi K., Luo L. (2014) Presynaptic partners of dorsal raphe serotonergic and GABAergic neurons. Neuron 83, 645–662.

Yan W. W., Xia M., Chiang J., Levitt A., Hawkins N., Kearney J., Swanson G. T., Chetkovich D., Nobis W. P. (2021) Enhanced Synaptic Transmission in the Extended Amygdala and Altered Excitability in an Extended Amygdala to Brainstem Circuit in a Dravet Syndrome Mouse Model. eNeuro 8.

Zeng C., Long X., Cotten J. F., Forman S. A., Solt K., Faingold C. L., Feng H.-J. (2015) Fluoxetine prevents respiratory arrest without enhancing ventilation in DBA/1 mice. Epilepsy Behav. 45, 1–7.

Zhang H., Zhao H., Zeng C., Van Dort C., Faingold C. L., Taylor N. E., Solt K., Feng H.-J. (2018) Optogenetic activation of 5-HT neurons in the dorsal raphe suppresses seizure-induced respiratory arrest and produces anticonvulsant effect in the DBA/1 mouse SUDEP model. Neurobiol. Dis. 110, 47–58.

Zhang R., Tan Z., Niu J., Feng H.-J. (2021) Adrenergic α2 receptors are implicated in seizure-induced respiratory arrest in DBA/1 mice. Life Sci., 119912.

